# Spatially Resolved Banff Tubulitis and Glomerulitis Scoring in Kidney Allograft Biopsies via Artificial Intelligent -Based Structure Segmentation and Spatial Transcriptomics

**DOI:** 10.64898/2026.05.08.723594

**Authors:** Heather R Kates, Chanel Lee, Anindya S. Paul, Ishaq Ansari, Anish Tatke, Tori Lee, Minh-Tri Nguyen, Michael T. Eadon, Pinaki Sarder, Yan Chen Wongworawat

**Author notes:** **Correspondences:** Yan Chen Wongworawat, MD, PhD, Department of Pathology and Human Anatomy, Loma Linda University Health, 11234 Anderson Street, Room 2124, Loma Linda, California 92354, USA, Pinaki Sarder, Ph.D., FASN, SMIEEE, SPIE Senior Member, Associate Professor, CMIL, Medicine - Quantitative Health, Department of Medicine, Division of Nephrology, University of Florida, 1600 SW Archer Road, Gainesville, FL 32610, USA. Co-last authors (Pinaki Sarder, Yan Chen Wongworawat).

## Abstract

**Background:** Tubulitis is a defining histologic feature of T cell-mediated rejection (TCMR), while glomerulitis is often characteristic of antibody mediated rejection (AMR). Histologic quantification of tubulitis and glomerulitis using Banff criteria is subject to interobserver variability. Bulk transcriptomic assays (e.g., MMDx) have introduced molecular correlations of tubulitis with TCMR and glomerulitis with AMR, but lack spatial resolution.

**Methods:** We applied a web-based platform, FUSION (Functional Unit State Identification in Whole Slide Images), to a cohort of 8 cases (n=2 per condition) with kidney allograft biopsy samples acute TCMR, active AMR, chronic active AMR, and no rejection (control). The machine-learning (ML) platform enabled integrated visualization and analysis of spatial transcriptomics (10x Genomics Visium v2) together with high-resolution whole-slide histology.

**Results:** Transcriptomics-derived immune cell proportions within AI-segmented tubular and glomerular regions were used to generate spatial Banff t- and g-scores. Derived t-scores showed full concordance with pathologist scores in both acute TCMR cases; g-scores showed concordance in 2 of 4 AMR cases, with discordant cases characterized by low absolute immune signal near the classification boundary.

**Conclusions:** We demonstrate the feasibility of using AI-based FTU segmentation integrated with spatial transcriptomics-derived immune cell proportions to generate spatially informed t- and g-scores aligned with Banff criteria, with full concordance in severe rejection and partial concordance in mild rejection. This approach lays the foundation for validated, spatial transcriptomics-augmented *t*-scores and *g*-scores that enhance diagnostic precision, reduces inter-observer variability among renal pathologists, and support potential clinical adoption.

## Introduction

Allograft biopsy remains the gold standard for diagnosis of kidney transplant rejection and is evaluated using the Banff classification, which defines antibody-mediated rejection (AMR) and T cell-mediated rejection (TCMR) based on specific histopathological and immunological criteria^1, 2^. AMR is classified as active, chronic active, or chronic, and requires evidence of acute tissue injury such as microvascular inflammation (glomerulitis and peritubular capillaritis), C4d positivity, donor-specific antibodies (DSAs), and chronic tissue injury such as transplant glomerulopathy. TCMR conversely is classified as acute or chronic active and is primarily determined by the extent of interstitial inflammation and tubulitis, although vascular injury may supersede in some cases^1, 2^.

According to the Banff criteria, tubulitis is graded by counting the number of mononuclear inflammatory cells within a tubular cross section or per ten tubular epithelial cells. On the other hand, glomerulitis is graded by the percentage of glomeruli involved. Histologic assessment is a time-consuming endeavor for nephropathologists, requiring careful annotation of immune and endothelial cells.

^3^In order to support nephropathologists in this burdensome task, digital pathology tools have been developed that use AI models to provide more objective and reproducible quantification of histological features. Enormous strides have been made by tools leveraging machine learning to allow pathologists to process complex visual data with high precision, detect subtle morphologic patterns, and achieve more quantitative and consistent evaluations than traditional manual assessment alone. However, image-based AI tools face inherent challenges in cellular localization, such as distinguishing intracytoplasmic inflammatory cells from epithelial nuclei in tubulitis assessment, or intraluminal leukocytes from endothelial nuclei in glomerulitis scoring. Transcriptomic data offers a complementary strategy by quantifying immune infiltration based on gene expression rather than solely on morphologic identification, and we hypothesized that integrating both approaches would enable us to detect these histologically complex lesions.

In this study, we used FUSION (Functional Unit State Identification in Whole Slide Images), an integrated visualization and analytic platform that combines histomorphological evaluation with spatial omics data on high-resolution whole slide images. By linking AI-based functional tissue unit (FTU) segmentation with Visium-derived cell-type deconvolution in FUSION, we performed compartment-specific quantification of immune cell proportions within individual tubules and glomeruli, allowing us to derive and visualize transcriptomically-informed tubulitis and glomerulitis scores in kidney allograft biopsies spanning the spectrum of rejection diagnoses.

## Materials and Methods

### Spatial Transcriptomics Data Processing

Spatial transcriptomics data were generated using the 10x Genomics Visium v2 platform (55 μm spot diameter) on FFPE kidney allograft biopsy sections from a cohort of 8 cases across four diagnostic groups: non-rejection control, active AMR, acute TCMR, and chronic active AMR. Four slides were processed, each containing two tissue sections. Raw data were processed using Space Ranger (10x Genomics). Additional details of this dataset have been published previously^4^. Spot-level quality control, filtering, and normalization were performed in R using Seurat (v5.1.0). Spots were filtered based on minimum transcript count (nCount ≥ 800) and feature count (nFeature ≥ 400).

### AI Segmentation and Cell Type Deconvolution

Anatomical segmentation of FTUs including tubules, non-sclerotic glomeruli, sclerotic glomeruli, cortical interstitium, vessels, and medullary interstitium was performed using FUSION (Functional Unit State Identification in Whole Slide Images), a web-based platform developed by the University of Florida Computational Microscopy Imaging Laboratory^5^. FUSION incorporates automated cell and FTU segmentation using the DeepCell Whole Slide Image pipeline, and cell-type deconvolution of Visium spots through the Seurat v5 function TransferData, using a comprehensive Kidney Precision Medicine Project (KPMP) single-nucleus atlas (>200,000 nuclei) as the reference to predict cell type proportions at each spot ^6-10^. Individual KPMP cell type labels were mapped to broader functional categories using FUSION-defined mapping; multiple immune cell types in the KPMP reference (including T cells, B cells, monocytes, macrophages, and other immune populations) were consolidated into a single immune category (IMM), with analogous groupings applied for proximal tubule (PT), loop of Henle (TAL/DTL/ATL), collecting duct (PC/CNT/IC), and other categories^9^. The immune cell proportion (IMM_prop) at each spot represents the aggregated deconvolved fraction of all IMM-mapped cell types relative to all cell types at that location. Aggregation of cell type proportions to the structure level were performed using FUSION’s FTU Aggregation plugin; aggregated annotations were exported from HistomicsUI as JSON files and parsed in R using jsonlite (v1.8.9) to extract per-FTU IMM proportions (“IMM_prop”) for downstream tubulitis and glomerulitis scoring.

### Tubulitis Scoring

Tubulitis was assessed for the acute TCMR slide and the non-rejection control slide. Per Banff convention, the t-score is determined by the most severely inflamed tubule in non-severely atrophic tubules (defined as having a diameter >=25% of that of unaffected or minimally affected tubules) in cortex (worst tubule criterion) ^2^. Accordingly, severely atrophic tubules are excluded from the IMM_prop calculation. In addition, because the tubulitis must be present in at least two foci, rare mildly inflamed tubules observed in non-rejection case 1 — located in a very focal inflammatory area near the corticomedullary junction — was manually excluded ^2^. For each tissue case, the maximum IMM_prop across all FTU-aggregated tubules was used as the scoring metric. Banff t-score thresholds were set a priori based on the correspondence between IMM_prop and expected cell counts: Visium spots (∼55µm diameter) capture approximately 5–15 cells, and tubules typically overlap one to two spots, such that an aggregated IMM_prop > 0.70 reflects immune cellular composition consistent with the Banff t3 criterion of >10 mononuclear cells per tubular cross-section; t2 (5–10 cells) and t1 (1–4 cells) thresholds were set at IMM_prop > 0.40 and > 0.15, respectively. Automated scoring was implemented in R and applied identically to both the TCMR and non-rejection control slides.

### Glomerulitis Scoring

Glomerulitis was assessed for the two AMR slides, active AMR and chronic active AMR, and the non-rejection control slide. Per Banff convention, the g-score is determined by the percentage of non-sclerotic glomeruli showing immune infiltration^2^. Globally sclerotic glomeruli were excluded from scoring by FUSION’s segmentation pipeline, which classifies sclerotic and non-sclerotic glomeruli as distinct FTU types. For each tissue case, glomeruli with at least one overlapping Visium spots were retrieved from the FUSION’s FTU Aggregation export and parsed in R using jsonlite (v1.8.9). A glomerulus was considered inflamed if its aggregated IMM_prop exceeded 0.10, a threshold selected to reflect the Banff criterion that even minimal immune infiltration is sufficient for classification as affected; unlike tubulitis, the Banff g-score is agnostic to infiltration severity within individual glomeruli and is determined entirely by the proportion affected^1, 2^. The percentage of inflamed glomeruli among those with Visium spot coverage was converted to a Banff *g*-score (g0: 0%; g1: >0–25%; g2: 25–75%; g3: >75%) and applied identically to all slides. Automated scoring was implemented in R.

## Results

### Demographics, Clinical, and Histopathologic Characteristics

We selected eight cases based on the histopathological and clinical features summarized in Table 1, representing four diagnostic groups: 1) non-rejection conditions, 2) active AMR, 3) acute TCMR, 4) chronic active AMR ^4^. Histopathologic diagnoses were established by our renal pathologists based on the 2018 Banff Criteria ^1, 2^.

**Table 1.** Study cohort, diagnostic condition, case identifiers, pathologist-assigned Banff scores and clinical features for all 8 cases. Adapted from Chen Wongworawat et al. Table 1 ^4^.

|  | Case 1 | Case 2 | Case 3 | Case 4 | Case 5 | Case 6 | Case 7 | Case 8 |
| --- | --- | --- | --- | --- | --- | --- | --- | --- |
| Diagnostic category | Non-rejection |  | Active AMR |  | Acute TCMR |  | Chronic active AMR |  |
| Pathologic diagnosis | Acute CNI toxicity | Very subtle ATI | C4d-positive active AMR with intimal arteritis | C4d-negative active AMR | Acute TCMR, grade 1B, plasma cell rich | Acute TCMR, grade 2A | Chronic active AMR with transplant arteriopathy | Chronic active AMR |
| Banff scores | t1, i0, v0, g0, ptc0, ci0, ct0, cg0, ti0, i-IFTA0, pvl0, C4d0 | t0, i0, v0, g0, ptc0, ci0, ct0, cg0, ti0, i-IFTA0, pvl0, C4d0 | t1, i2, v1, g2, ptc2, ci0, ct0, cg0, ti2, i-IFTA0, pvl0, C4d3 | t0, i0, v0, g1-2, ptc2, ci0, ct0, cg0, ti0, i-IFTA0, pvl0, C4d1 | t3, i3, v0, g0, ptc0, ci0, ct0, cg0, ti3, i-IFTA0, pvl0, C4d1 | t3, i2, v1, g0, ptc2, ci0, ct0, cg0, ti2, i-IFTA0, pvl0, C4d1-2 | t0, i0, v0, g1-2, ptc0, ci0, ct0, cg1b, ti0, i-IFTA0, pvl0, C4d2 | t0, i0, v0, g1, ptc1, ci0, ct0, cg2, ti0, i-IFTA0, pvl0, C4d2 |
| Age | 41 | 58 | 53 | 31 | 28 | 36 | 41 | 49 |
| Cause of ESKD | Hepatorenal syndrome | Diabetes | Unknown | Hypoplastic kidney | Unknown | Unknown | Unknown | Unknown |
| SCr (mg/dL) | 1.5 | 1.9 | 6.1 | 1.5 | 18 | 10 | 1.36 | 1.4 |
| DSA | Positive | Positive | Positive | Positive | Positive | Positive | Negative | Positive |
| Graft function | DGF | DGF | Normal | Normal | Normal | Normal | Normal | Normal |

CNI, acute calcineurin inhibitors; AMR, active antibody mediated rejection; ATI, toxicity acute tubular injury; Banff Score: tubulitis (t), interstitial inflammation in non-scarred areas (i), intimal arteritis (v), glomerulitis (g), peritubular capillaritis (ptc), interstitial fibrosis (ci), tubular atrophy (ct), glomerular basement membrane double contours (cg), total inflammation (ti), inflammation in the area of IFTA (i-IFTA), polyomavirus load (pvl); DGF, delayed graft function; DSA, donor specific antibody; ESKD, end-stage kidney disease; IFTA, interstitial fibrosis and tubular atrophy; Scr, Serum creatine; TCMR, cell mediated rejection.

### Tubulitis is consistently identified within spatially resolved tubular regions and correlates with the Banff t score

Tubulitis was assessed across 943 FTU-aggregated tubules from the acute TCMR slide (*n* = 498 and *n* = 445 tubules for Cases 1 and 2, respectively) and 684 tubules from the non-rejection control slide (*n* = 463 and *n* = 221 for Cases 1 and 2). The highest tubule IMM proportion was used as the scoring metric in accordance with the Banff worst-tubule criterion. Both acute TCMR cases achieved full concordance with pathologist t-scores: Case 1 (*i* ≥ 2 and *t*3, which meets criteria for Banff grade 1B) and Case 2 (*v1* regardless of *i* or *t*, which meets criteria for Banff grade 2A) both yielded worst-tubule IMM proportions of 0.856 and 0.886, respectively, exceeding the *a priori* t3 threshold (IMM_prop > 0.70), with 10.0% and 5.8% of tubules above the t1 threshold (Table 2, Figure 1). Qualitative visualization of tubular IMM proportions on the H&E showed the highest-IMM tubules corresponding to regions of apparent tubular injury in both TCMR cases (Figure 1).

**Table 2.**
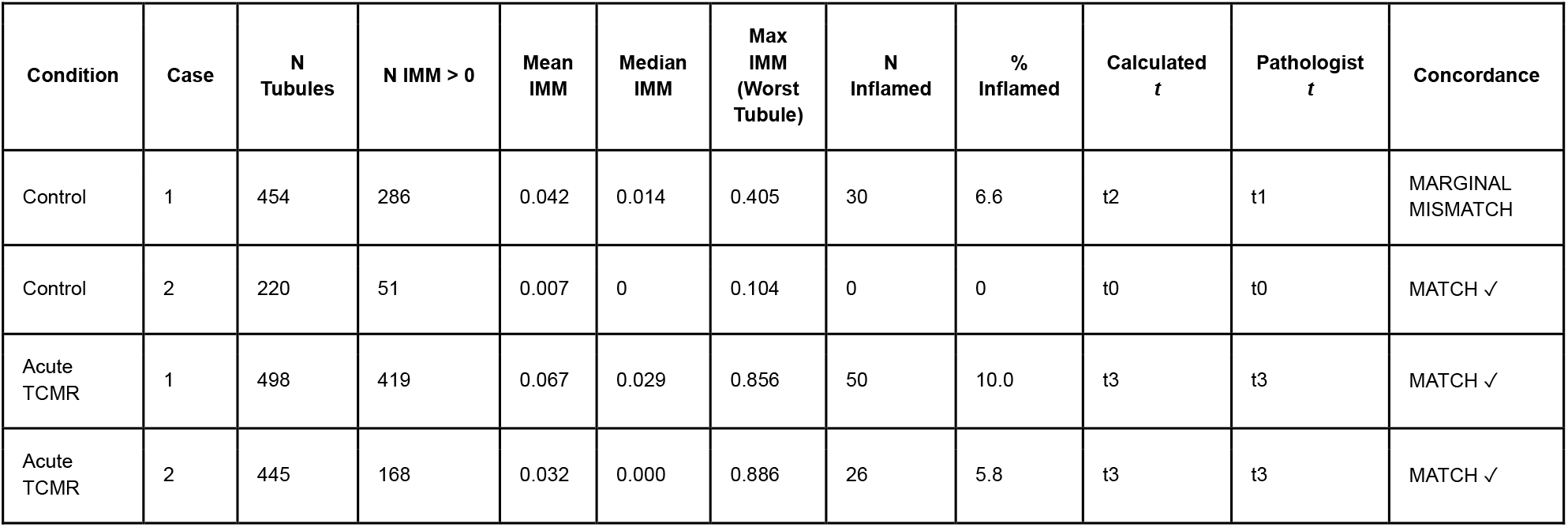
Tubulitis scoring results. Per-case summary of FTU-aggregated tubule-level IMM proportions and calculated Banff t-scores for acute TCMR and non-rejection control cases. N Tubules: total tubules with ≥1 Visium spot; Max IMM: worst-tubule IMM proportion used as the scoring metric per Banff worst-tubule criterion; % Inflamed: percentage of tubules exceeding the t1 threshold (provided for descriptive purposes only; t-score is determined by the worst single tubule per Banff convention); Calculated t: Banff t-score derived from fixed thresholds (t1 > 0.15, t2 > 0.40, t3 > 0.70); Pathologist t: score assigned by expert renal pathologist; Concordance: match/mismatch with pathologist score.

**Figure 1.**
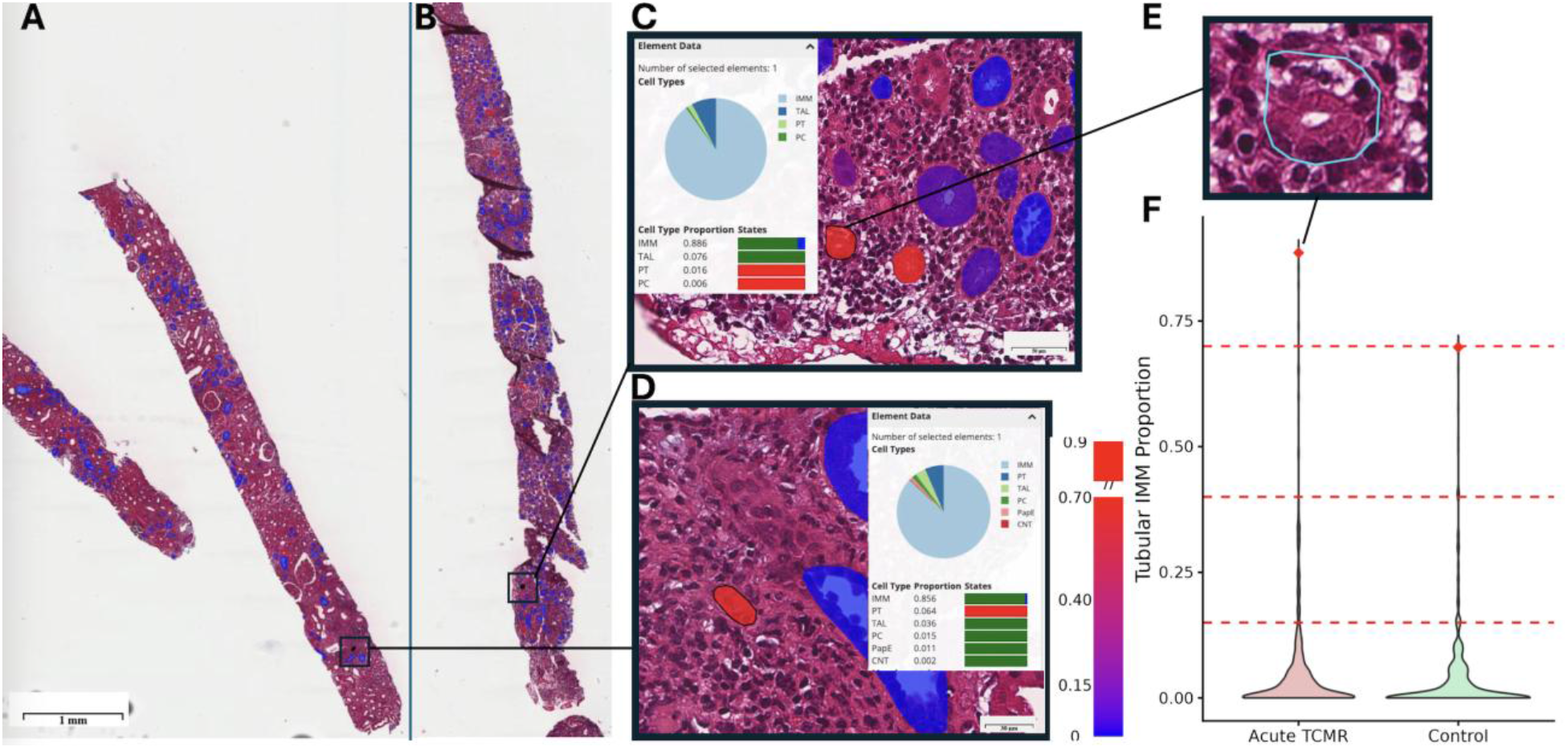
Automated tubulitis scoring in acute TCMR. (A) Acute TCMR whole-slide H&E at 2.3× magnification without Visium spots; scale bar = 1 mm. (B) Same slide with Visium spots colored by tubular IMM proportion (blue–red, 0–0.70; color bar shown); worst tubule outlined in black. **(C)** High-magnification inset of the worst tubule in Case 2 (IMM proportion = 0.886) with FUSION cell type proportions shown; IMM constitutes 88.6% of deconvolved signal. **(D)** Further zoomed inset of the worst tubule in Case 1 (IMM proportion = 0.856) with FUSION cell type proportions shown; black outline indicates the worst tubule; IMM constitutes 85.6% of deconvolved signal. **(E)** Maximum magnification inset (∼10 μm tubule) of the worst tubule from Case 1, outlined in cyan, demonstrating dense mononuclear infiltration of a single small tubular cross-section; this tubule corresponds to the red diamond in panel F. **(F)** Violin plots showing the distribution of tubular IMM proportions for acute TCMR and non-rejection control; red diamond indicates the worst tubule per condition; dashed red lines indicate Banff t-score thresholds (t1 > 0.15, t2 > 0.40, t3 > 0.70). Visium spots shown in semi-transparent yellow.

Application of the same fixed thresholds to the non-rejection control slide produced broadly concordant scores for both control cases against pathologist scores of t1 and t0, although the worst-tubule IMM proportion for Case 1 (0.405) marginally exceeded the t2 threshold (IMM_prop > 0.40) we classified its calculated *t-*score as t1 due to known variability in spot-to-tubule assignment at tubule boundaries. Worst-tubule IMM proportions in control tissue (0.405 and 0.104) were substantially lower than in TCMR (Table 2, Figure 1).

### Glomerulitis quantified within spatially resolved glomerular regions showed partial concordance with pathologist Banff g-scores

Glomerulitis was assessed across glomeruli with Visium spot coverage from the active AMR slide (*n* = 4 and *n* = 14 glomeruli for Cases 1 and 2), the chronic active AMR slide (*n* = 10 and *n* = 4 for Cases 1 and 2), and the non-rejection control slide (*n* = 10 for Case 2; Case 1 had no glomeruli overlapping Visium spots and could not be scored). A glomerulus was classified as inflamed if its aggregated IMM proportion exceeded 0.10, consistent with the Banff criterion that even minimal immune infiltration is sufficient for classification as affected. The percentage of inflamed glomeruli was converted to a Banff g-score (Table 3).

**Table 3.** Glomerulitis scoring results. Per-case summary of FTU-aggregated glomerulus-level IMM proportions and calculated Banff g-scores for active AMR, chronic active AMR, and non-rejection control cases. N Glom: total non-sclerotic glomeruli with ≥1 Visium spot (Note: Case 1 had no glomeruli overlapping Visium spots and could not be scored); % Affected: percentage of glomeruli with IMM_prop > 0.10; Calculated g: Banff g-score; Pathologist g: score assigned by expert renal pathologist; Concordance: match/mismatch with pathologist score.

| Condition | Case | N Glom | N IMM > 0 | Max IMM | N Affected | % Affected | Calculated $g$ | Pathologist $g$ | Concordance |
| --- | --- | --- | --- | --- | --- | --- | --- | --- | --- |
| Control | 2 | 10 | 9 | 0.039 | 0 | 0 | g0 | g0 | MATCH ✓ |
| Active AMR | 1 | 4 | 4 | 0.619 | 4 | 100 | g3 | g2 | MISMATCH |
| Active AMR | 2 | 14 | 13 | 0.041 | 0 | 0 | g0 | g1-2 | MISMATCH |
| Chronic Active AMR | 1 | 10 | 10 | 0.170 | 3 | 30 | g2 | g1-2 | MATCH ✓ |
| Chronic Active AMR | 2 | 4 | 3 | 0.023 | 0 | 0 | g0 | g1 | MISMATCH |

Two of four AMR cases achieved concordance with pathologist g-scores. Active AMR Case 1 scored g3 (4/4 glomeruli affected, 100%) against a pathologist score of g2; the discrepancy is attributable to the small denominator; with only 4 glomeruli captured over Visium spots, any affected glomerulus yields ≥25% and all 4 affected yields 100%, precluding a g2 calculation regardless of threshold. Chronic Active AMR Case 1 correctly scored g2 (3/10 glomeruli affected, 30%), within the pathologist’s range of g1-2. The two discordant cases (Active AMR Case 2: max IMM_prop 0.041) and Chronic Active AMR Case 2: max IMM_prop 0.023) had no glomeruli exceeding the 0.10 threshold, with immune signal well below the classification boundary. Qualitative visualization of glomerular IMM proportions overlaid on H&E showed that the highest-IMM glomeruli in concordant cases corresponded to regions of apparent immune cell enrichment, while glomeruli in discordant cases showed no appreciable immune signal (Figure 2). The non-rejection control correctly scored g0 for Case 2, with a maximum glomerular IMM proportion of 0.039 (Table 3, Figure 2).

**Figure 2.**
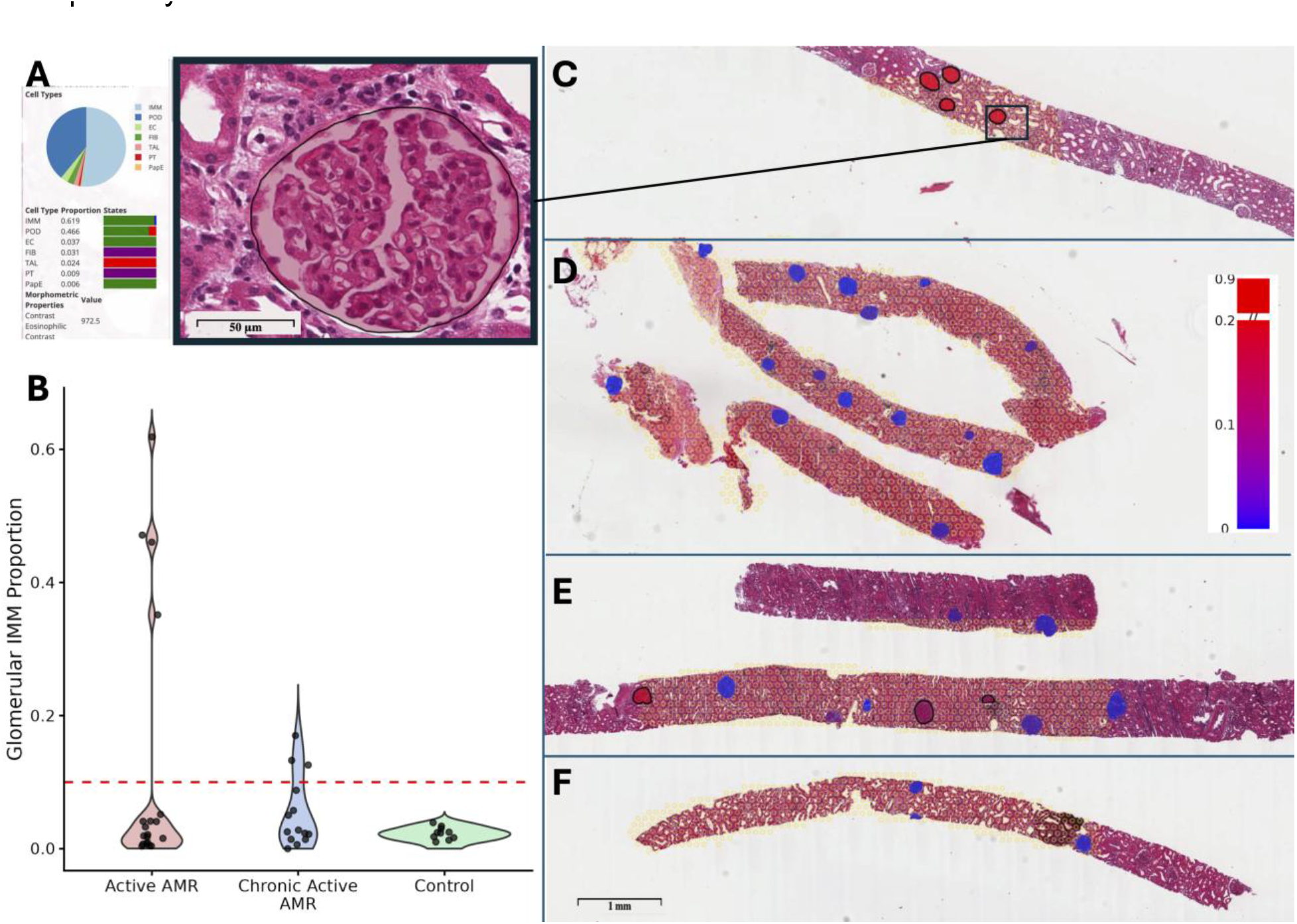
Automated glomerulitis scoring across diagnostic conditions. **(A)** Representative high-magnification inset of the highest-IMM glomerulus in Active AMR Case 1 (IMM proportion = 0.619), with FUSION cell type proportions shown; black outline indicates IMM proportion > 0.10. **(B)** Violin plots showing the distribution of glomerular IMM proportions for Active AMR, chronic active AMR, and non-rejection control; individual glomeruli overlaid as points; dashed red line indicates the glomerulitis threshold (IMM proportion = 0.10). **(C, D)** Active AMR Cases 1 and 2: non-sclerotic glomerular FTU polygons colored by IMM proportion (blue–red, 0–0.20; color bar shown) overlaid on H&E at 1.4× magnification; scale bar = 1 mm; glomeruli with IMM proportion > 0.10 outlined in black. Case 1: 4/4 glomeruli affected (calculated g3, pathologist g2). Case 2: 0/14 glomeruli affected (max IMM proportion 0.041, calculated g0, pathologist g1-2). **(E, F)** Chronic Active AMR Cases 1 and 2. Case 1: 3/10 glomeruli affected (calculated g2, pathologist g1-2, concordant). Case 2: 0/4 glomeruli affected (max IMM proportion 0.023, calculated g0, pathologist g1, discordant). Visium spots shown in semi-transparent yellow.

## Discussion

This study presents a proof-of-concept for automated, spatially resolved Banff scoring of tubulitis and glomerulitis in kidney allograft biopsies, integrating AI-based FTU segmentation in FUSION with cell type deconvolution from the Visium spatial transcriptomics platform. Using AI-based FTU segmentation and immune cell proportion (IMM) thresholds derived from the expected correspondence between spot-level deconvolved cell fractions and Banff cell count criteria, we achieved full concordance with pathologist t-scores in both acute TCMR cases and partial concordance with pathologist g-scores in AMR. To our knowledge, this represents the first application of spatially resolved transcriptomics to automate structure-level Banff scoring in kidney allograft rejection. As a proof-of-concept study with n=2 cases per diagnostic category, this work is designed to establish methodological feasibility rather than validate diagnostic performance. Validation in larger, prospectively collected cohorts will enable formal sensitivity and specificity estimation with meaningful confidence intervals. Cases achieving concordance between pathologists and spatially derived scores were characterized by high absolute immune signal. Both TCMR cases exhibited worst tubule IMM proportions exceeding 0.85, and the concordant AMR case had 30% of captured glomeruli above the inflammation threshold. In cases where rejection is severe and immune infiltration is widespread, the correspondence between deconvolved IMM proportion and pathologist-observed infiltration was robust. Discordant cases were those with mild pathologist scores and low absolute immune signal, with maximum glomerular IMM proportions below 0.05 in both discordant AMR cases.

Several technical characteristics of the Visium V2 assay limit detection of mild immune infiltration. The 55 μm spot diameter is large relative to both individual tubules and glomerular capillary loops, such that a spot overlapping with a structure boundary captures signal averaged over both the target FTU and surrounding interstitium, diluting the apparent IMM proportion when infiltration is focal or mild. For tubulitis, where tubules typically overlap only one to two spots, the worst-tubule metric is often derived from a single Visium spot and is sensitive to severe diffuse infiltration but vulnerable to noise at lower severity. The marginal control tubulitis mismatch (worst-tubule IMM 0.405, exceeding the t2 threshold by a negligible margin, against a pathologist score of t1 likely reflect real but diagnostically insignificant low-level immune signal rather than true rejection, particularly in the setting of minimal cortical interstitial inflammation (i0).

Beyond spot-size limitations, IMM proportions are deconvolved estimates derived by label transfer from a reference atlas rather than direct cell counts, and may not reflect true immune cell abundance, which is especially limiting near the classification boundary where small errors in deconvolved proportions could determine whether a structure is called inflamed or not. Finally, the tissue area assessed by the pathologist is larger than the Visium capture area, which covers only a fraction of the biopsy core and does not guarantee spot overlap with every structure. This disproportionately affects mild rejection, where the probability of capturing the single worst-affected structure decreases as the fraction of affected structures decreases.

Although Banff criteria require immune cells to be present within specific anatomical compartments (within the tubular epithelium for tubulitis, and within glomerular capillary lumina for glomerulitis), spot-level IMM proportions reflect aggregated signal from all cells within the 55 μm spot footprint regardless of their precise location within the FTU. Therefore, this approach cannot distinguish intraepithelial and peritubular signal nor capillary luminal from mesangial or interstitial signal. Subcellular resolution platforms such as Xenium *in situ*, which can localize individual transcripts to specific cellular compartments, may be better suited to recapitulate the precise anatomical requirements of Banff scoring in future work.

In summary, this work demonstrates that spatially resolved transcriptomics integrated with AI-based FTU segmentation can recapitulate Banff tubulitis and glomerulitis scoring with full concordance in severe rejection and partial concordance in mild rejection, with discordant cases attributable to interpretable technical limitations. Validation in larger, more diverse cohorts, together with higher-resolution spatial platforms and continued advancements in computational deconvolution and lesion mapping, may further improve clinical translation and expand therapeutic strategies. While manual Banff scoring remains the clinical reference standard and is subject to interobserver variability, these results suggest that spatially resolved transcriptomics can complement expert review, though cost and technical requirements are likely to limit near-term adoption in routine practice.

## Author Contributions

Y.C.W conceived the research and supervised all aspects of the work. H.R.K, A.P, A.T, I.A and Y.C.W contributed to data analysis. H.R.K, Y.C.W, A.P, C.L and T.L drafted the manuscript. P.S supervised the research and laboratory activities. M.T.E and M.N served as consultants. All the authors read, edited, and approved the manuscript.

## Acknowledgments

We appreciate Histology Laboratory and California Tumor Tissue Registry, Department of Pathology and Human Anatomy at Loma Linda University, Loma Linda, California, for their assistance with FFPE tissue processing.

## Data Availability Statement

All analysis code is publicly available at https://github.com/SarderLab/LLU-kidney-spatial-tx. The data that supports the findings of this study are available on request from the corresponding author upon reasonable request.

